# Interferon Regulatory Factor 6 Determines Intestinal Epithelial Cell Development and Immunity

**DOI:** 10.1101/2024.01.04.574257

**Authors:** Austin P. Wright, Sydney Harris, Shelby Madden, Bryan Ramirez Reyes, Ethan Mulamula, Alexis Gibson, Isabella Rauch, David A. Constant, Timothy J. Nice

**Affiliations:** Department of Molecular Microbiology and Immunology, Oregon Health and Science University, Portland, OR 97239, USA

## Abstract

Intestinal epithelial cell (IEC) responses to interferon (IFN) favor antiviral defense with minimal cytotoxicity, but IEC-specific factors that regulate these responses remain poorly understood. Interferon regulatory factors (IRFs) are a family of nine related transcription factors, and IRF6 is preferentially expressed by epithelial cells, but its roles in IEC immunity are unknown. In this study, CRISPR screens found that *Irf6* deficiency enhanced IFN-stimulated antiviral responses in transformed mouse IECs but not macrophages. Furthermore, KO of *Irf6* in IEC organoids resulted in profound changes to homeostasis and immunity gene expression. *Irf6* KO organoids grew more slowly, and single-cell RNA sequencing indicated reduced expression of genes in epithelial differentiation and immunity pathways. IFN-stimulated gene expression was also significantly different in *Irf6* KO organoids, with increased expression of stress and apoptosis-associated genes. Functionally, the transcriptional changes in *Irf6* KO organoids were associated with increased cytotoxicity upon IFN treatment or inflammasome activation. These data indicate a previously unappreciated role for IRF6 in IEC biology, including regulation of epithelial development and moderation of innate immune responses to minimize cytotoxicity and maintain barrier function.

## Introduction

The interferon (IFN) family of cytokines are a first line of defense against viral pathogens. Activation of IFN receptors initiates a signaling pathway resulting in transcription of IFN-stimulated genes (ISGs), which include many direct-acting antiviral effectors (Schneider et al., 2014; Schoggins, 2019). There are three types of IFN, which are defined by their use of distinct membrane-bound receptors (Sadler & Williams, 2008). The transcriptional profiles (ISGs) induced by each IFN type overlap substantially, but there are cell type-specific differences in antiviral protection. Type I IFN can act on nearly every nucleated cell in the body, but type III IFN (IFN-λ) primarily acts on epithelial cells of barrier tissues, and intestinal epithelial cells (IECs) preferentially respond to IFN-λ (Baldridge et al., 2017; Mahlakoiv et al., 2015; Mordstein et al., 2010; Nice et al., 2015; Pott et al., 2011; Sommereyns et al., 2008; Van Winkle et al., 2022). For example, interferon lambda receptor KO (*Ifnlr1*^-/-^) mice fail to control intestinal replication of murine norovirus (MNV) (Nice et al., 2015), and homeostatic antiviral responses in the intestinal epithelium are absent in *Ifnlr1*^-/-^ mice (Van Winkle et al., 2022). Thus, understanding the factors that regulate IFN-λ responsiveness of IECs is of particular importance to intestinal health.

Type I and III IFN receptors can utilize the same canonical signaling pathway (Sadler & Williams, 2008). Receptor-associated Janus kinase 1 (JAK1) and tyrosine kinase 2 (TYK2) phosphorylate signal transducer and activator of transcription 1 (STAT1) and STAT2. Interferon regulatory factor 9 (IRF9) joins with STAT1 and STAT2 to form IFN-stimulated gene factor 3 (ISGF3), which translocates to the nucleus and binds IFN-sensitive response element (ISRE) motifs in ISG promoters (Sadler & Williams, 2008). One major difference between type I and III IFNs is the strength of signaling, with type III IFN resulting in a more moderate but sustained level of gene expression (Forero et al., 2019; Mendoza et al., 2017; Pervolaraki et al., 2018). Thus, a modest response stimulated by type III IFN in epithelial cells benefits tissue homeostasis by maintaining antiviral protection with minimal epithelial cytotoxicity (Van Winkle et al., 2020). However, IEC-specific factors that regulate the IFN response remain poorly understood.

IEC-specific regulators of the interferon response may include relatives of canonical signaling facters, such as members of the JAK, STAT, and IRF families. There are nine IRF proteins that share a conserved N-terminal DNA binding domain (DBD) that interacts with a conserved GAAA consensus DNA sequence that is part of the ISRE motif (Negishi et al., 2018; Taniguchi et al., 2001). The C-terminal regions are more divergent, and include regulatory motifs. IRFs 3-9 encode an IRF-association domain (IAD) and an autoinhibitory region that facilitate dimeric interaction and inhibition of dimerization, respectively. Despite the discovery of IRFs as regulators of IFN, and their homology in the DBD, some IRFs have been shown to regulate development of specific cell types. For example, IRF4 and IRF8 regulate leukocyte development (Gabriele & Ozato, 2007; Mancino & Natoli, 2016; Tsujimura et al., 2002; H. Wang et al., 2008), and IRF6 regulates keratinocyte development (Ingraham et al., 2006; Kousa et al., 2017; Kwa et al., 2015). IRF6 is expressed by all epithelial lineages, but developmental and immunological roles in the intestine were unknown.

To identify the presence of IEC-specific factors that regulate the antiviral IFN response, we designed a CRISPR screen that targeted canonical IFN signaling factors and homologous family members. We found that *Irf6* KO enhanced IFN-stimulated antiviral immunity of IEC cell lines but not macrophages. RNAseq analysis of *Irf6* KO IEC cell lines revealed substantial baseline changes in growth and development pathway genes, and dysregulated ISG expression that correlated with antiviral protection. We found that *Irf6* was highly expressed in primary IEC organoids and intestinal tissues. *Irf6* KO in primary IEC organoids reduced growth and developmental gene expression, with enhanced production of particular ISGs and increased IFN-stimulated cytotoxicity. These data suggest a previously unappreciated role for IRF6 in IEC development and immunity.

## Results

### Protection against virus-triggered death by IFN treatments in macrophage and epithelial cell lines

To study genetics of IFN-stimulated antiviral protection, we used BV2 and M2C-CD300lf cell lines that represent myeloid-lineage and intestinal epithelial-lineage, respectively. First, we directly compared the efficacy of type I and III IFNs in these cells by performing dose-response titrations using recombinant murine IFN-β and IFN-λ (**Fig. 1**). Both BV2 and M2C cells were treated with IFN for 24 hours before being challenged with a lytic strain of murine norovirus (MNV) (Robinson et al., 2019; Van Winkle et al., 2018). The BV2 cells were infected at an MOI=10 resulting in <1% viability, and the M2C cells were infected at an MOI=50 resulting in ∼9% viability. IFN-β treatment protected MNV-infected M2C from death, reaching ∼100% viability at 10 ng/mL (**Fig. 1A**, squares). IFN-λ treatment also protected MNV-infected M2C from death, but with a lower maximum survival rate of ∼30% (**Fig. 1A**, circles). The BV2 macrophages had a survival rate of ∼10% when treated with 10ng/ml IFN-β, but showed no increase in survival when pretreated with any dose of IFN-λ, as expected (**Fig. 1B**). To compare differences in responsiveness between IFN type and cell type in the subsequent CRISPR screen, we selected doses that moderately increased viability following MNV infection: **1)** 10ng/ml IFN-λ-treated M2C IECs, **2)** 0.01 ng/ml IFN-β-treated M2C IECs, and **3)** 10ng/ml IFN-β-treated BV2 macrophages (**Fig. 1**, dashed lines).

**Figure 1.**
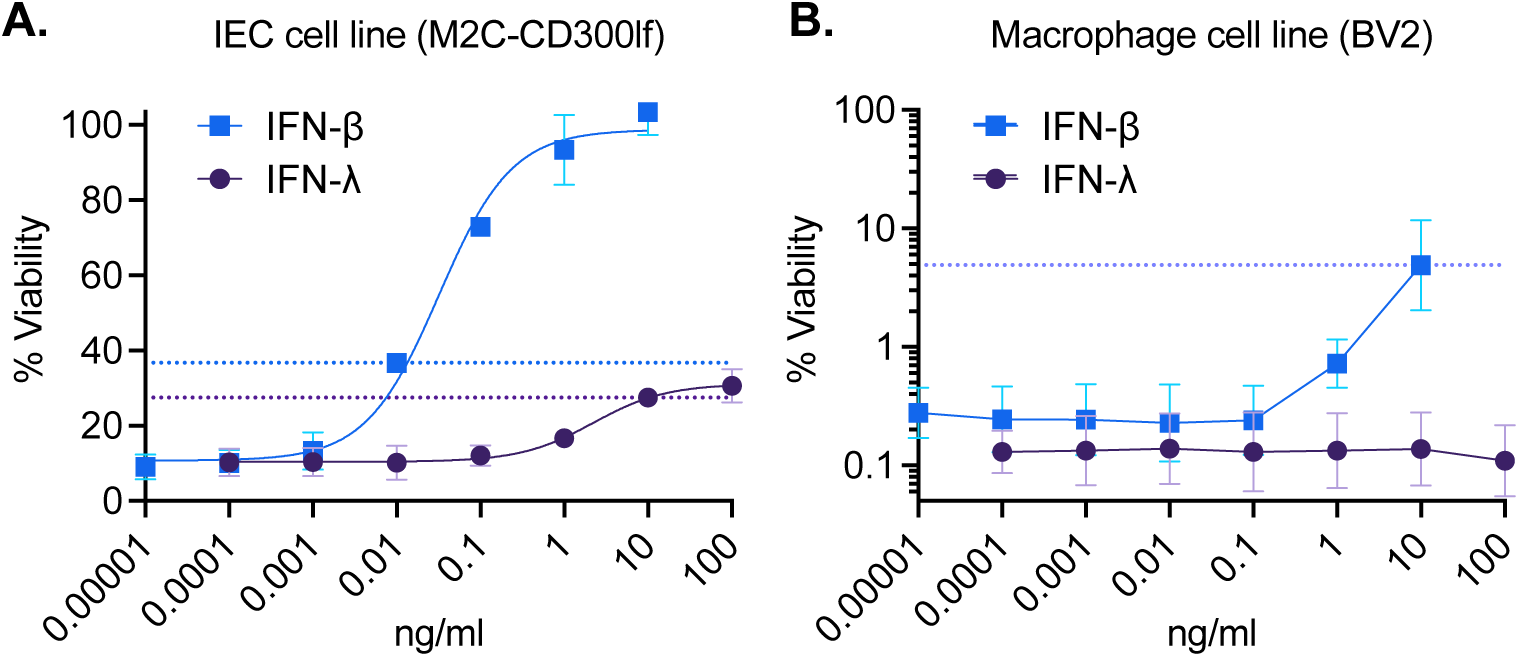
Protection against norovirus-triggered death by IFN treatments in macrophage and epithelial cell lines. Dose-response curves with IFN-β and IFN-λ pre-treatment of **A**) M2C-CD300lf epithelial cells and **B**) BV2 macrophage cells. ATPglo viability for each dose was normalized as a percent of uninfected, untreated cells. Dashed lines indicate doses selected for use in subsequent screens. Data is represented as mean and standard deviation of two (M2C) or three (BV2) replicates.

### CRISPR screens for IEC-specific regulators of the IFN response

To determine requirement of candidate genes for IFN-stimulated protection, we knocked out genes within JAK, STAT, NF-κB and IRF families using CRISPR lentivirus transduction (two gRNAs/gene, **Table S1**). For each IFN treatment and cell type, we saw that gRNA targeting of canonical signaling factors resulted in lower protection provided by IFN treatments, validating our screening approach (**Fig. 2**). In particular, IFN-β treatment of BV2 cells with gRNA targeting *Stat1, Stat2, Irf9,* and *Jak1* resulted nearly no protection (0.01-1%), whereas treatment of non-targeting controls resulted in 3-11% protection (**Fig. 2A-B**). gRNA targeting *Irf1* and *Tyk2* may have a more modest effect on IFN-stimulated protection of BV2 cells, resulting in an intermediate amount of protection (1-3%) following IFN-β treatment (**Fig. 2A-B**). None of the CRISPR targeted BV2 cell lines showed increased IFN-β-stimulated protection relative to controls.

**Figure 2.**
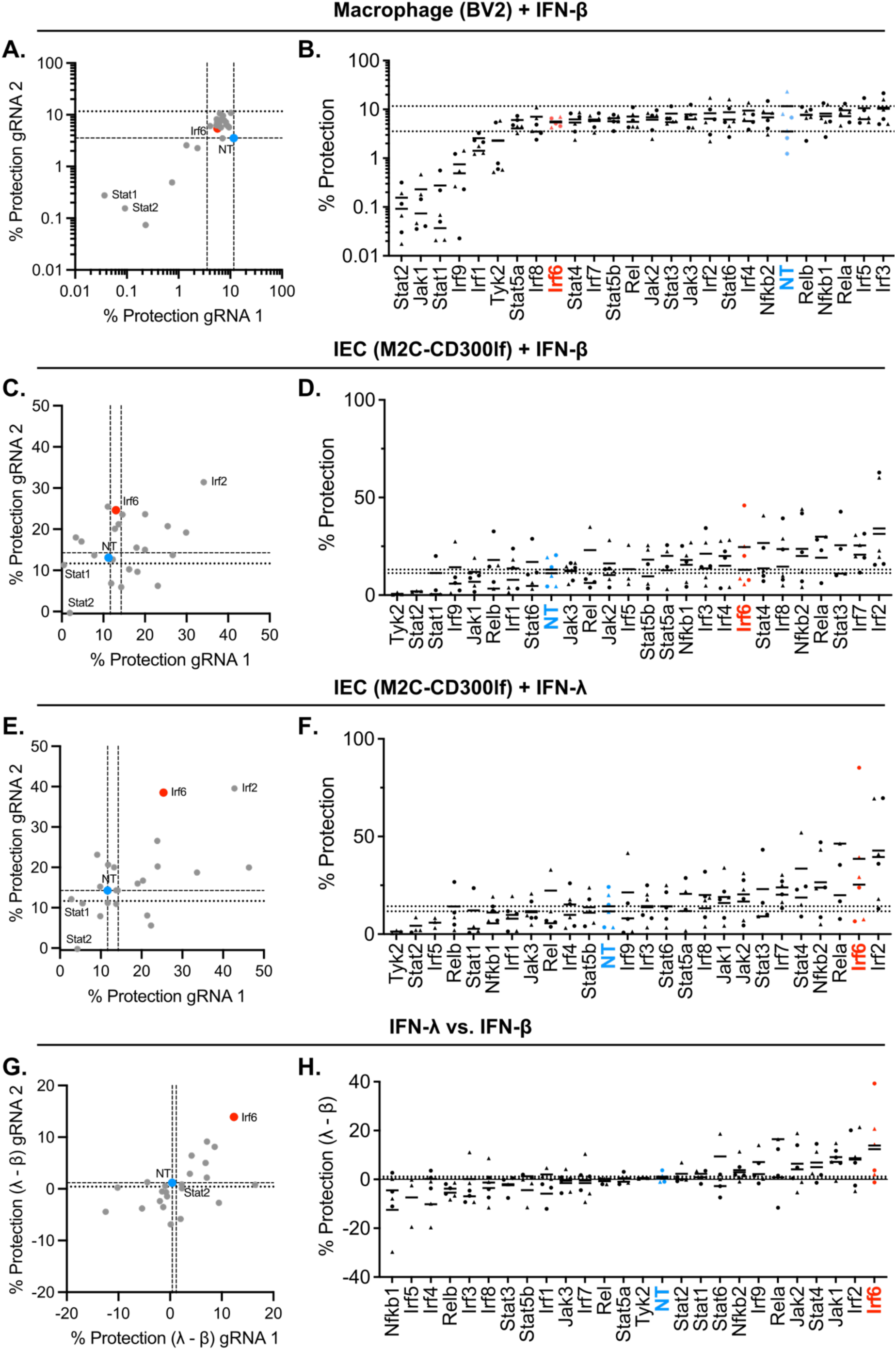
CRISPR screen for IFN-stimulated protection of macrophage and IEC cell lines. CRISPR KO cells were screened for differences of IFN-stimulated protection from MNV. IFN-stimulated protection was calculated by subtracting viability of untreated controls from IFN-treated cells (% protection, **A-F**), and differences between IFN types were determined by subtracting IFN-β-stimulated protection from IFN-λ-stimulated protection (λ – β, **G-H**). Data is plotted as individual replicates (**B, D, F, H**) or as the mean values from three replicate experiments for each of two independent gRNAs per gene (**A, C, E, G**). **A-B.** IFN-β-stimulated protection of BV2 macrophages. **C-D.** IFN-β-stimulated protection of M2C-CD300lf IECs. **E-F.** IFN-λ -stimulated protection of M2C-CD300lf IECs. **G-H.** Difference between IFN-λ- and IFN-β-stimulated protection of M2C-CD300lf IECs. Genes positioned in the bottom left quadrant of **G** are more protected by IFN-β than IFN-λ and genes positioned in the upper right quadrant are more protected by IFN-λ than IFN-β. Dotted lines in all plots represent the mean values of non-targeting control gRNAs (blue). Shapes for individual replicate datapoints represent each independent gRNA. Mean values are indicated for each gRNA. Data represents three experimental replicates.

Similar to BV2 cells, M2C-CD300lf cells with gRNA targeting *Stat1* and *Stat2* were among the least protected cells following IFN-β treatment (**Fig. 2C-D**). Likewise, IFN-λ treatment of M2C-CD300lf cells with gRNA targeting of *Stat1* and *Stat2* resulted in reduced protection relative to non-targeting controls (**Fig. 2E-F**). However, unlike the BV2 cells, there were several genes where gRNA targeting increased the IFN-stimulated protection of M2C-CD300lf cells, including *Irf2* and *Irf6*. Targeting of *Irf2* resulted in increased protection of M2C-CD300lf cells pretreated with either IFN-β or IFN-λ (**Fig. 2C-F**), indicating that *Irf2* may inhibit IFN signaling in these epithelial cells. This is consistent with previously described inhibitory activity of *Irf2* (Harada et al., 1989). Targeting of *Irf6* resulted in increased protection of M2C-CD300lf cells pretreated with IFN-λ (**Fig. 2E-F**), but appeared to have a more modest or inconsistent effect on M2C-CD300lf cells pretreated with IFN-β (**Fig. 2C-D**), and had no effect on BV2 macrophages (**Fig. 2A-B**).

To quantify differences between IFN-λ and IFN-β treatments, we determined the difference in IFN-stimulated protection for each CRISPR gRNA in M2C-CD300lf cells (**Fig. 2G-H**). Notably, this comparison was between IFN types within the same cell lines, thereby minimizing effects of variation in MNV susceptibility between cell lines (**Table S2**). We found that *Irf6*-targeted cells had the largest difference between IFN-λ and IFN-β treatments, with greater protection provided by IFN-λ than IFN-β (**Fig 2G-H**). These results suggested that *Irf6* is a novel regulator of the IFN-stimulated antiviral response in IECs.

To increase confidence in selecting candidate genes for further study, we complemented the viability CRISPR screen with an orthogonal FACS-based pooled CRISPR screen (**Fig. 3A**). Pooled CRISPR-transduced cells were pre-treated with IFN types, infected with MNV, and cells with the greatest production of MNV protein (top 10% NS1/2-positive) were sorted for quantification of gRNA abundance (**Fig. 3A**). MNV NS1/2 protein staining at 8 hours post-infection was consistently detected in the BV2 and M2C-CD300lf pools (**Fig. 3B-C**, ‘no IFN’ group), indicating that these cell lines are similarly capable of supporting MNV replication. IFN-β treatment resulted in lower fluorescence intensity of MNV NS1/2 in both BV2 and M2C cells (**Fig. 3B-C**). However, IFN-λ treatment of M2C cells did not significantly reduce MNV NS1/2 protein staining, confounding our ability to identify genes that influence IFN-λ activity by this screening method. Notably, these data suggest that actions of IFN-λ to protect cells from MNV-triggered death (**Fig. 1-2**) are distinct from those that block viral protein production (**Fig. 3B-C**). Therefore, IFN-λ-treated cells were not further considered in analysis of this pooled CRISPR screen.

**Figure 3.**
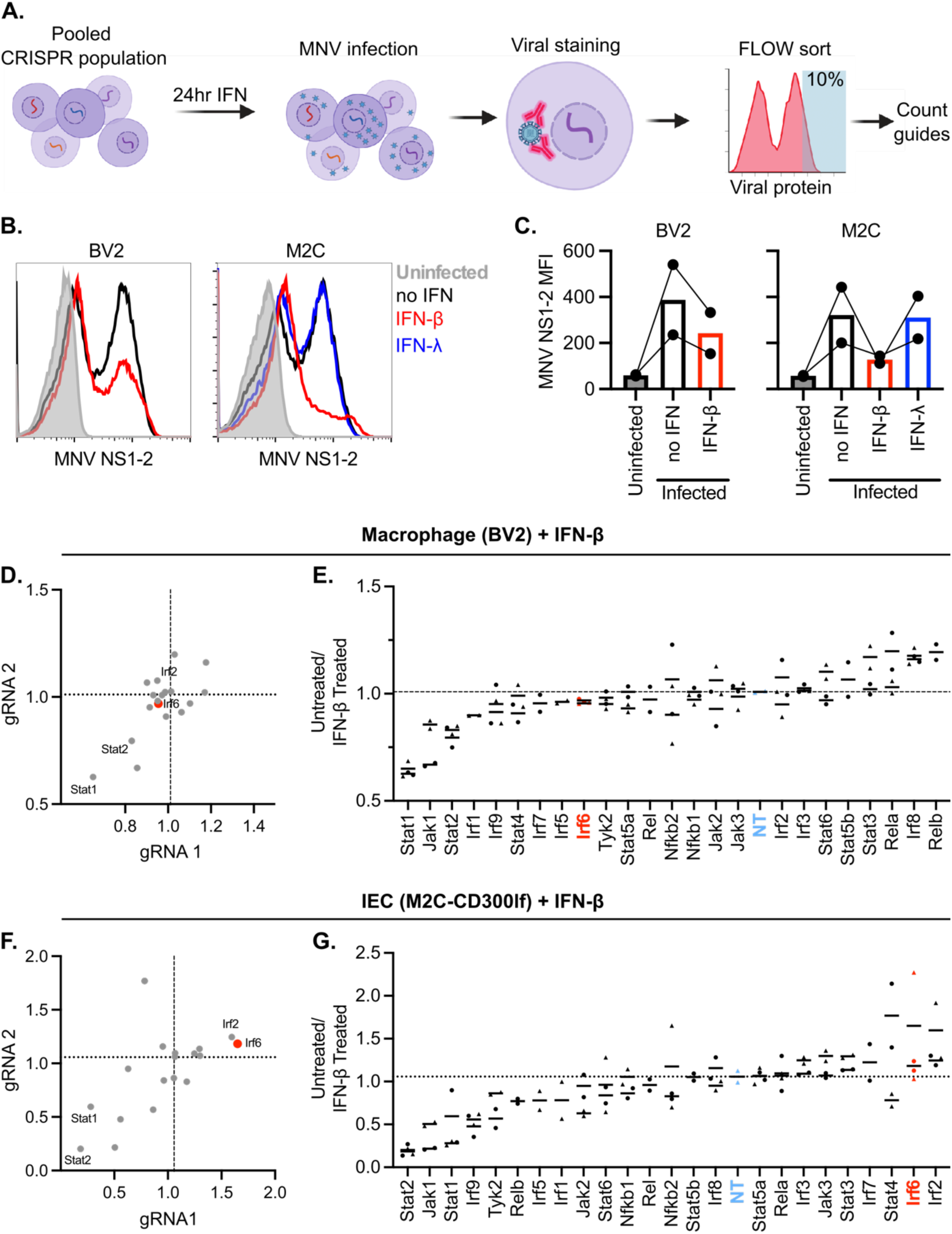
Pooled CRISPR screen for IFN-stimulated antiviral response in macrophage and IEC cell lines. **A.** Pooled CRISPR transduced cells were screened for genes that altered IFN-stimulated protection from MNV infection by cell sorting the top 10% of infected cells based on staining for MNV NS1-2 protein production. **B.** Representative FACS plots of NS1-2 staining. **C.** Mean fluorescence intensity (MFI) of MNV NS1-2 staining 8 hours post-infection of cells pre-treated for 24 hours with no IFN, 1 ng/ml IFN-β, or 100 ng/ml IFN-λ, as indicated. Each dot represents the mean fluorescence intensity of a single replicate. **D-G.** Plotted values indicate abundance of each gRNA in untreated cells divided by abundance of the same gRNA in cells pretreated with IFN-β. Dashed line indicates mean of non-targeting control. **D, F.** Mean values of the two gRNAs for each gene plotted on x and y axes. **E, G.** Plotted values of each replicate, with the gRNAs for each gene represented as distinct symbols. Mean values for each gRNA are indicated. Genes are ranked from left to right in order of enhancement to inhibition of the IFN response. Data represents two experimental replicates.

We sequenced gRNAs present within the top 10% of MNV NS1/2-positive cells, and compared gRNA counts between groups. Genes that promote IFN-stimulated antiviral immunity were expected to have correspondingly decreased gRNA counts within untreated groups relative to IFN-treated groups (**Table S3**). Indeed, canonical genes (*Stat1*, *Stat2*, *Jak1*) were decreased within untreated M2C and BV2 cells relative to paired IFN-β-treated groups, whereas non-targeting control gRNAs were equally represented (**Fig. 3D-G**). These expected outcomes validate our screen results. Analogous to the results of the viability screen (**Fig. 2C-D**), *Irf2* was increased within untreated M2C relative to the paired IFN-β-treated groups (**Fig. 3F-G**), but was not different in BV2 cells (**Fig. 3D-E**). Likewise, *Irf6* was increased within untreated M2C relative to the paired IFN-β-treated groups (**Fig. 3F-G**), analogous to the results from IFN-λ-treated cells in the viability screen (**Fig. 2E-F**). Thus, both CRISPR screening approaches suggested a novel and cell type-specific role for *Irf6* in the regulation of IFN-stimulated antiviral response of IECs.

### *Irf6* KO slows growth and alters IFN-stimulated protection of an IEC cell line

Both CRISPR screens suggested that targeting of *Irf6* resulted in greater IFN-stimulated antiviral protection of M2C IECs. To further test the role of *Irf6*, we generated monoclonal cell lines targeted by the two *Irf6* gRNAs used in the screen, and sequence-verified disruption of the *Irf6* locus. *Irf6* gRNA 1 cut directly before the conserved DNA binding domain, and *Irf6* gRNA 2 cut near the beginning of the predicted IRF-association domain (**Fig. 4A**). We selected monoclonal cell lines with mutations that resulted in frame shift and early stop codons (**Fig. 4A**). *Irf6* qPCR from the KO cell lines indicated undetectable (gRNA 1, KO1) or significantly reduced (gRNA 2, KO2) *Irf6* mRNA expression (**Fig. 4B**). Notably, the baseline abundance of *Irf6* mRNA was low in all M2C cells (greater than 1000-fold less abundant than the housekeeping gene *Rps29*, **Fig. 4B**), and we were unable to detect Irf6 protein by western blot. However, we observed phenotypic alterations following clonal isolation of the *Irf6* KO cells, with fewer cells harvested during expansion compared to non-targeting controls, and several instances of large and multinucleated cells within *Irf6* KO2 isolates (**Fig. 4C**). These observations suggested monoclonal isolates may be selected for adaptation to *Irf6* deficiency. To quantitate the growth phenotype, we counted cells over time after plating and found that that both *Irf6* KO M2C cell lines had a decreased growth rate, with significantly fewer cells recovered compared to non-targeting controls (**Fig. 4D**).

**Figure 4.**
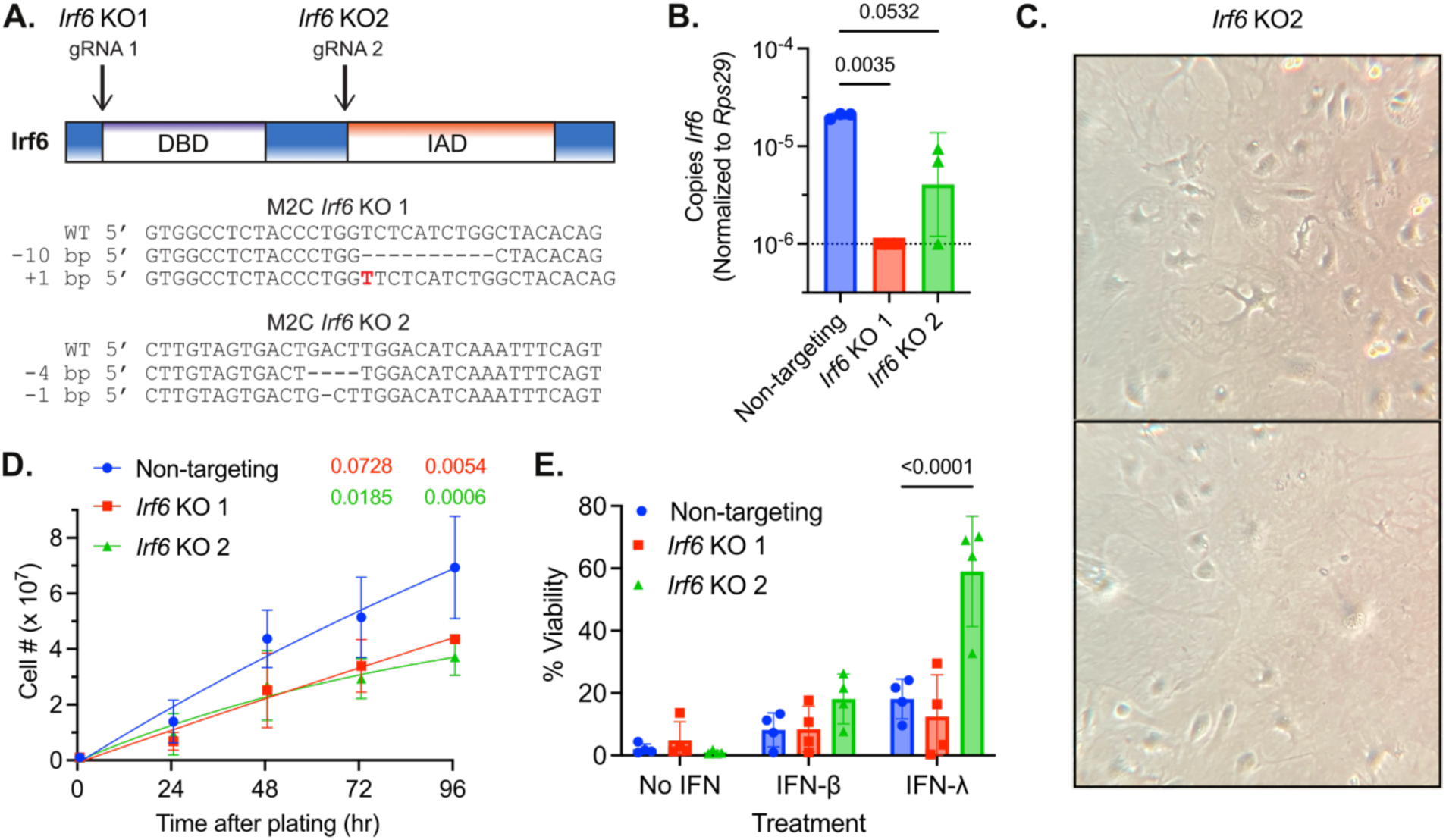
*Irf6* KO slows growth and alters IFN-stimulated protection in an IEC cell line. **A.** Graphical representation of targeting sites for each *Irf6* gRNA in the context of resulting protein domains, and sequences of monoclonal cell lines selected for further study. DBD = DNA-binding domain. IAD = IRF-association domain. *Irf6* KO1 had a 10bp deletion and a 1bp insert resulting in early stop codons. *Irf6* KO2 had a 4bp deletion and a 1bp deletion resulting in early stop codons. **B.** qPCR of *Irf6* of three replicates. Dashed line indicates limit of detection. P-values calculated by one-way ANOVA. **C.** Representative images of large multinucleated *Irf6* KO M2Cs. **D.** Growth curves from three experimental replicates. P-values are shown above for 72 and 96 hour timepoints, calculated by two-way ANOVA. **E.** ATPglo viabiliy assay 24 hours after MNV infection of cells pre-treated for 24 hours with no IFN, 0.01 ng/mL IFN-β, or 10 ng/mL IFN-λ. Data points represent four experimental replicates. P-values calculated by two-way ANOVA.

We tested IFN-stimulated protection of KO cells by measuring viability after MNV infection, with or without IFN pretreatments, as in the original CRISPR screen (**Fig. 2**). To ensure uniform susceptibility of monoclonal isolates to MNV infection, we re-transduced them with lentivirus encoding CD300lf. The resulting cell lines were equally susceptible to MNV, with less than 10% surviving in the absence of IFN pretreatment (**Fig. 4E**). IFN-β and IFN-λ pretreatment increased viability of all cell lines after MNV infection. There were no significant differences in IFN-β-stimulated protection between the cell lines, but *Irf6* KO2 had modest increase (two-fold) in average viability after IFN-β pre-treatment. In contrast, there were significant differences in IFN-λ-stimulated protection between the cell lines, with *Irf6* KO2 increased to 60% average viability after IFN-λ treatment, compared to 20% average viability in control cells (**Fig. 4E**). *Irf6* KO1 did not have significantly different viability from controls after either type of IFN pretreatment, but exhibited high variance across the four replicates, with lower IFN-stimulated protection in two experiments (**Fig. 4E**). Thus, there was an inconsistent effect of *Irf6* KO on IFN-stimulated protection from MNV in M2C-CD300lf cells. However, there was a consistent reduction in growth rate for both *Irf6* KO, suggesting that Irf6 plays an important homeostatic role in these cells.

### *Irf6* KO alters baseline and IFN-stimulated gene expression

To better understand the baseline and IFN-stimulated growth and viability phenotypes, we performed RNA sequencing on the *Irf6* KO and control cell lines. We harvested RNA from cells plated and treated in parallel to the viability assay in **figure 4E**, including both untreated and IFN-treated groups. Principal component analysis of the RNA sequencing results clustered groups with primary separation based on cell identity (PC1, 50% variance) and secondary separation based on IFN treatment (PC2, 20% variance) (**Fig. S1A**). Consistent with PCA analysis, there were hundreds of significantly different genes at baseline in *Irf6* KO cell lines compared to non-targeting controls (**Table S4**). In *Irf6* KO1 we saw 103 up-regulated differentially expressed genes (DEGs), and 247 down-regulated DEGs; in *Irf6* KO2 we saw 1274 up-regulated DEGs and 1088 down regulated DEGs; both *Irf6* KO cell lines shared 31 up-regulated DEGs and 93 down-regulated DEGs (**Fig. S1B**). There was a notable cluster of DEGs that were uniformly down-regulated in both *Irf6* KO cell lines (**Fig. S1C**, box), and additional baseline DEGs that were unique to KO2. Pathway analysis of DEGs for each *Irf6* KO cell line revealed shared significant changes in pathways that regulate cell differentiation and growth (**Fig. S1D**). These enriched pathways are consistent with the decreased growth rate of these *Irf6* KO cell lines (**Fig. 4D**). Several genes decreased in both *Irf6* KO cell lines are part of the “cell differentiation” GO pathway, including fibroblast growth factor 13 (*Fgf13*), Fms related receptor tyrosine kinase 4 (*Flt4*), *Notch1*, Lymphoid enhancer binding factor 1 (*Lef1*), Microtubule-associated protein 2 (*Map2*), and Slit guidance ligand 2 (*Slit2*) (**Fig. S1E**). Notably, previous ChIP-seq experiments from human keratinocytes (Botti et al., 2011) identified IRF6-bound loci in 12 human orthologs of genes down-regulated at baseline in both *Irf6* KO M2C cell lines (*SLIT2*, *CELSR1*, *SMARCA1*, *CAMK1D*, *PEG10*, *KPNA3*, *LTBP1*, *JAM2*, *KDR*, *AIG1*, *PCDH17*, and *EIF4G3*). These data indicate substantial changes to baseline gene expression in *Irf6* KO IEC cell lines, and suggest roles for Irf6 in growth and differentiation of IECs.

*Irf6* KO2 M2C cells had additional changes in baseline gene expression beyond the growth and development genes shared between KO cell lines (**Fig. S1A-E**). Pathway analysis of *Irf6* KO2 DEGs indicated significant enrichment of genes in “immune system process”, “inflammatory response”, and “regulation of leukocyte migration” GO pathways (**Fig. S1D**, **Table S4**). These immune-related genes upregulated at baseline in *Irf6* KO2 included *Cxcl10*, *Tnip3*, *Lbp*, *Ikbkg*, *Tnfrsf9*, *Il33*, *Ccl2*, *Ccl20*, and *Ifnlr1* (**Fig. S1E**). The enrichment of *Ifnlr1* is particularly notable because it correlates with the increased IFN-λ-stimulated viability seen in **figure 4E**.

To compare the expression of IFN-regulated genes in these cell lines, we compared IFN-stimulated samples to replicate untreated controls (**Fig. S1F-G**). A heat map of all IFN-regulated genes reveals that most of them are upregulated by IFN treatments, an expected characteristic of ISGs (**Fig. S1F, Table S4**). To compare the magnitude of IFN responsiveness, we plotted the Log2 fold-change for the IFN-regulated genes. These comparisons show significantly higher stimulation by IFN-λ in *Irf6* KO2 compared to non-targeting controls, and significantly less stimulation of *Irf6* KO1 by both IFN-β and IFN-λ (**Fig. S1G**). The overall increase in ISGs seen in *Irf6* KO2 treated with IFN-λ correlates with the increased viability following MNV infection (**Fig. 4E**).

Taken together, these RNAseq data indicate a consistent down-regulation of growth and differentiation genes in *Irf6* KO cell lines. In contrast, immunity-related genes and ISGs exhibit divergent phenotypes between the *Irf6* KO cell lines that may reflect unique adaptations to *Irf6* deficiency. Although the primary goal of our CRISPR screen was to identify novel regulators of the IFN response in IECs, these data suggest that Irf6 plays more foundational roles in IEC biology at baseline.

### *Irf6* is expressed in primary IECs and regulates organoid homeostasis

Transformed cell lines such as M2C may selectively downregulate IRF6 due to its role as a tumor suppressor (Bailey & Hendrix, 2008; Botti et al., 2011; Restivo et al., 2011). Indeed, we saw that *Irf6* was minimally expressed in M2C cells (**Fig. 4B**, **6A**). So, we sought to test *Irf6* expression and function in primary cells. We found that *Irf6* expression in mouse small intestine and colon tissues was >10,000-fold higher than the M2C cell line, and *Irf6* in spleen tissue was significantly lower than intestinal tissues (**Fig. 5A**). Primary IEC organoids derived from mouse small intestine expressed *Irf6* at levels within the same order of magnitude as intestinal tissues (**Fig. 5A**). These results indicate that the M2C IEC cell line expresses sub-physiological levels of *Irf6*, so we focused subsequent study of Irf6 on primary IECs.

**Figure 5.**
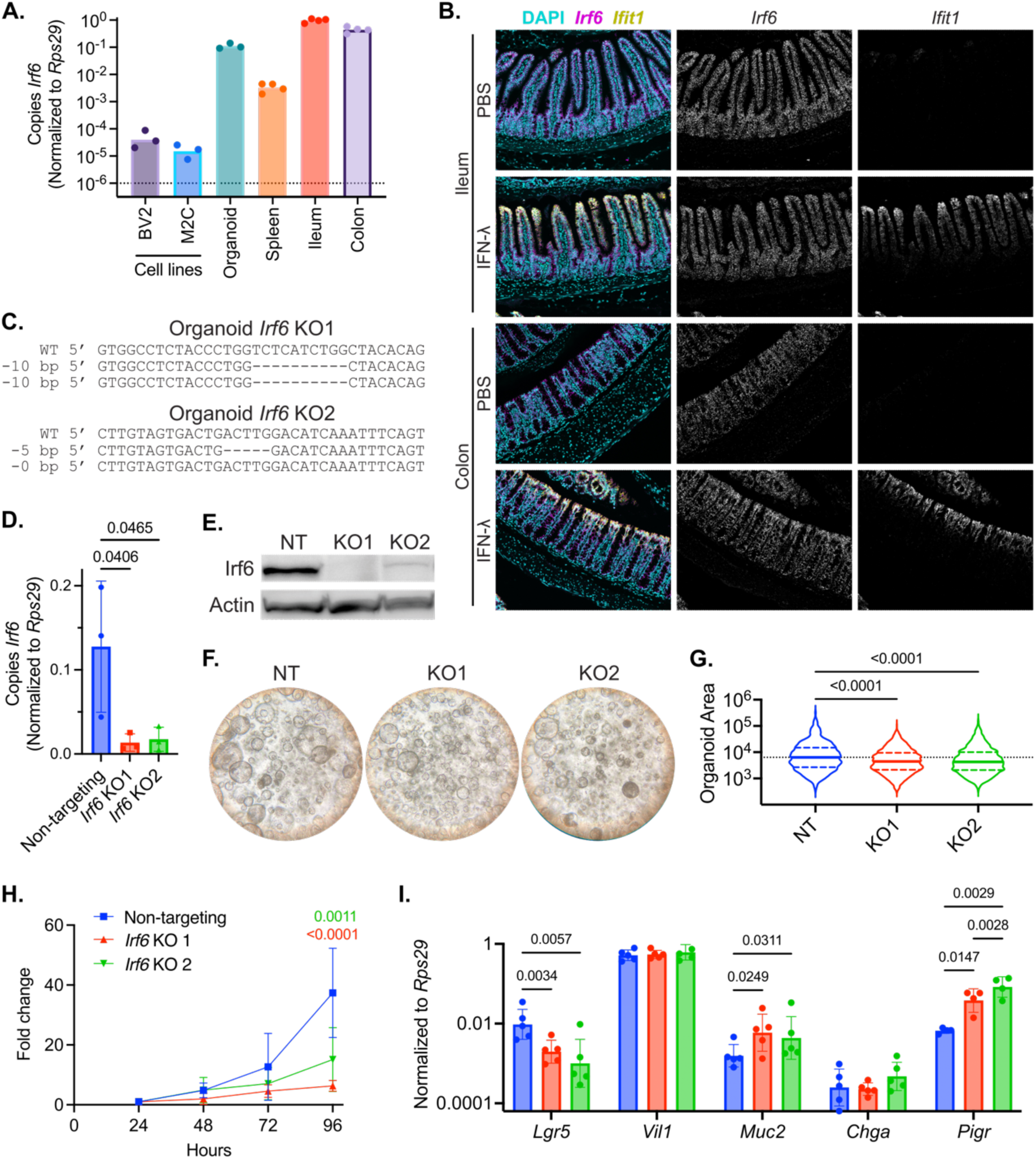
*Irf6* is expressed in primary IECs and regulates organoid homeostasis. A. Expression of *Irf6* in different cells and tissues. **B.** *Irf6* and *Ifit1* (ISG) expression in small intestine and colon of adult mice. Treatment with PBS or 3 μg peg-IFN-λ, as indicated, 24hr prior to tissue collection. Representative of four mice per group. **C.** Sequence of *Irf6* locus in monoclonal IEC organoid lines transduced with CRISPR lentivirus. **D.** *Irf6* expression by qPCR. **E.** Irf6 protein expression by western blot in the non-targeting CRISPR control organoids (NT) compared to *Irf6* KO organoids, representative of three replicates. **F.** Representative photos of organoids two days after plating. **G.** Cross-sectional area of organoids measured by ImageJ (arbitrary units) two days after plating. Violin plots show the median and quartiles of three experimental repeats (n = 1242, 1756, 1483). Dashed line indicates median of non-targeting control. P-values calculated by Kruskal-Wallis test. **H.** Growth curves of IEC organoids from three experimental replicates, normalized within each replicate to cell number at 24 hours. **I.** Expression of indicated genes by qPCR in *Irf6* KO organoids and non-targeting controls. Data points represent five experimental replicates. P-values calculated by two-way ANOVA.

**Figure 6.**
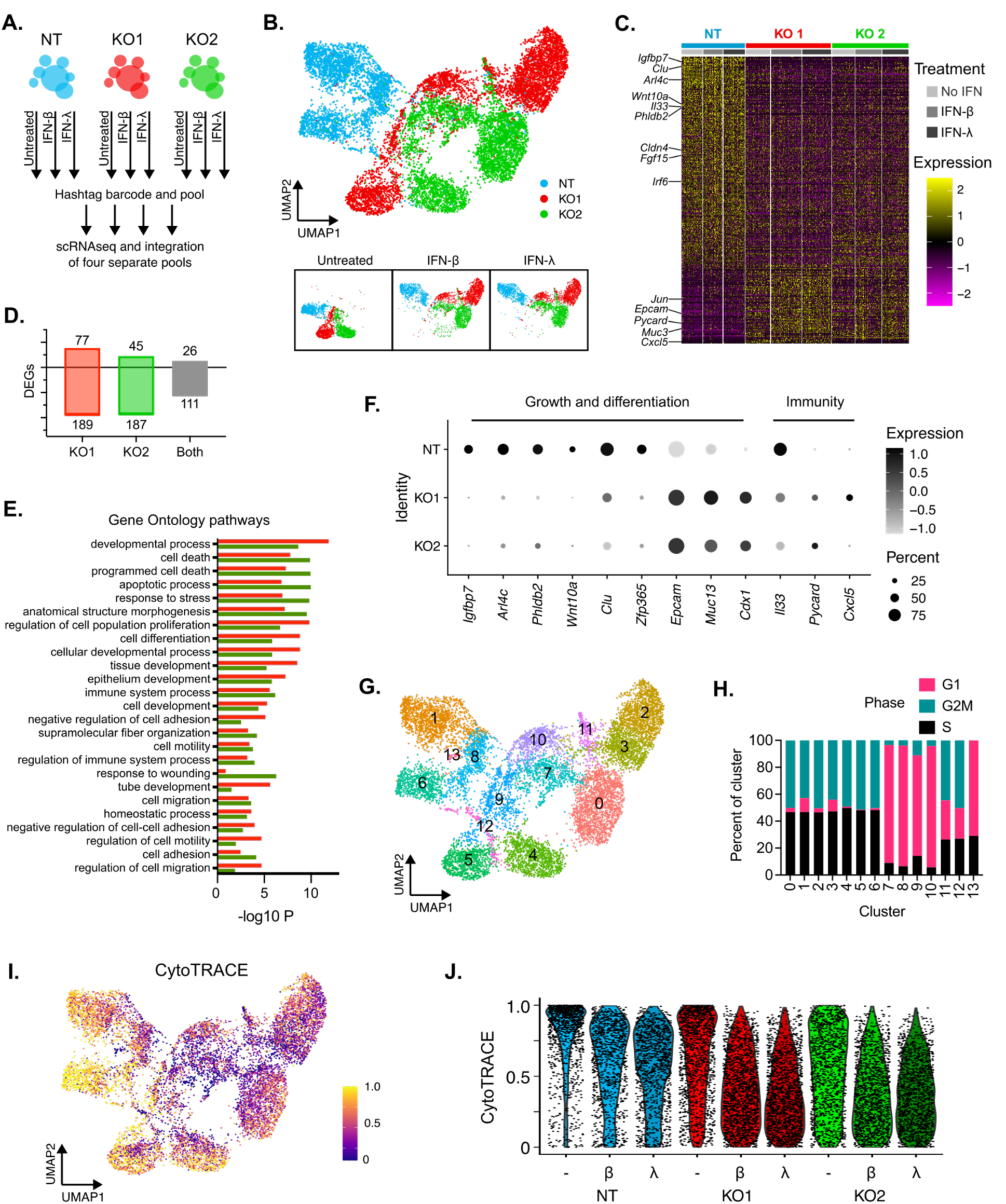
*Irf6* regulates development and immune response genes in primary IECs. Non-targeting (NT) or *Irf6* KO organoids were treated with no IFN, 10 ng/mL IFN-β, or 25 ng/mL IFN-λ for 24 hours prior to preparation of single cells for scRNA-seq. **A.** Diagram depicting experimental groups, multiplexing, and pooling strategy. Two pools consisted of organoid lines transduced with the MNV receptor CD300lf, with or without MNV infection. **B.** UMAP multidimensional clustering of all sequenced cells, colored by cell line. Insets at bottom are split by IFN treatment group, as indicated. **C.** Heatmap of *Irf6*-dependent DEGs arranged by preferential expression in NT cells (top) to preferential expression in *Irf6* KOs (bottom). Selected genes related to development and immunity are labeled. **D.** Number of DEGs from **C** for each KO organoid line compared to NT control, and overlapping DEGs shared by KO organoid lines. **E.** Association between genes in **C-D** and selected GO pathways for *Irf6* KO1 (red bars) and *Irf6* KO2 (green bars) organoid lines. **F.** Feature plots depicting distribution of selected genes shared with other IRF6 datasets or GO pathways, as indicated by headings. **G.** Identification of clusters with distinct gene expression profiles among all groups (**B**). **H.** Distribution of cell cycle phase categories within each cluster. **I-J.** CytoTRACE analysis of differentiation. Higher CytoTRACE score indicates more stem-like cells. All analyses performed on integrated data from four single-cell pools (**A**). DEGs were defined as >1.5-fold change with adjusted p-value <0.05 using analysis pipelines described in methods.

To visualize distribution of Irf6 within intestinal tissues we performed *in situ* hybridization for *Irf6* in ileum and colon of mice injected with PBS or IFN-λ 24 hours prior to tissue collection (**Fig. 5B**). In all intestinal tissues, *Irf6* was predominant within the epithelium, and was similar in abundance from crypt base to mature enterocytes and colonocytes (**Fig. 5B**). The ISG response in mice injected with IFN-λ was assessed by detection of *Ifit1*, and expression of this ISG was predominant within mature IECs (**Fig. 5B**). *Irf6* transcripts were not strikingly different between IFN-λ-treated mice and PBS controls, suggesting that *Irf6* is not an ISG (**Fig. 5B**). These imaging analyses indicated that *Irf6* is expressed in IECs of small intestine and colon, including IFN-λ-responsive cells.

We next sought to generate *Irf6* KO primary IECs by transducing organoids with *Irf6*-targeting CRISPR lentiviruses used in the screens (**Fig. 4A**). We selected transduced IEC organoid clones and sequenced gRNA target sites within the *Irf6* locus to assess gene disruption. We identified a homozygous *Irf6* KO in organoids transduced with CRISPR gRNA 1 (**Fig. 5C**, KO1). However, in organoids transduced with CRISPR gRNA 2 (cuts after DBD), we recovered only clones with heterozygous targeting of the *Irf6* gene (**Fig. 5C**, KO2). Analysis of *Irf6* expression by qPCR showed significant decreases in both KO organoid lines (**Fig. 5D**). Western blot of Irf6 showed no detectable protein in KO1 organoids and a substantially decreased protein level in KO2 organoids (**Fig. 5E**). We speculated that homozygous targeting of *Irf6* using gRNA 2 may be more deleterious due to potential expression of a protein fragment containing the Irf6 DBD only (**Fig. 4A**). We were unable to visualize any such fragment on western blot, but a similar heterozygous deletion has been linked to a human orofacial clefting syndrome (Degen et al., 2020), indicating the potential for biological activity of this heterozygous truncation. So, we included both *Irf6* KO organoid lines in our subsequent studies.

During culture of *Irf6* KO IEC organoids, we noticed that they appeared smaller and darker (**Fig. 5F**). To quantify organoid size, we took pictures two days after plating and measured the cross-sectional area of organoids in each image. *Irf6* KO organoids were significantly smaller than the non-targeting control organoids (**Fig. 5G**). Additionally, counting cells over time post-plating revealed that *Irf6* KO organoids grew significantly more slowly than non-targeting controls (**Fig. 5H**). Thus, *Irf6* deficiency reproducibly results in slower growth of primary IECs, similar to the baseline phenotype observed in *Irf6-*deficient M2C cell lines.

The slower growth rate of *Irf6* KO organoids, together with the earlier observation of decreased growth and development genes in *Irf6* KO M2C cell lines (**Fig. S1D-E**), led us to investigate the expression of IEC differentiation genes in organoids. *Lgr5* is expressed by intestinal stem cells, and is reduced in expression as enterocytes mature; *Lgr5* was 5- to 10-fold lower in *Irf6* KO organoids relative to non-targeting controls (**Fig. 5I**). *Vil1* is an enterocyte marker, and was not significantly different in *Irf6* KO organoids. *Muc2* is a mucin glycoprotein produced by goblet cells; we saw a ∼5-fold increase in *Irf6* KO organoids relative to non-targeting controls (**Fig. 5I**). *Chga* is marker for enteroendocrine cells; *Chga* was not significantly different in *Irf6* KO organoids. *Pigr* is an Fc-receptor that facilitates translocation of immunoglobulin A into the intestinal lumen, and is stimulated by innate responses to microbiota; *Pigr* was 5- to 10-fold higher in the *Irf6* KO organoids (**Fig. 5I**). Taken together, these data indicate that *Irf6* deficiency in primary IEC organoids results in slower growth, reduced size, and increased expression of certain differentiation genes.

### *Irf6* regulates development and immune response genes in primary IECs

To define *Irf6*-dependent alterations to IEC organoid gene expression, we prepared single-cell RNA sequencing libraries from four pools of organoid cells, with or without IFN treatments (**Fig. 6A, methods**). Upon demultiplexing and integration of these four single-cell pools, we ended up with 12,151 cells suitable for analysis. Dimensional reduction of the integrated single-cell data revealed that a primary source of variation (UMAP1) was related to *Irf6* KO and a secondary source of variation (UMAP2) was due to IFN treatments (**Fig. 6B**). Separate clustering of the untreated groups confirmed that a primary source of variation was related to *Irf6* KO, independent of IFN treatment (**Fig. S2A**).

To identify global *Irf6*-dependent transcriptional changes, we compared gene expression in each *Irf6* KO organoid line to non-targeting controls. Hundreds of genes were significantly different in each *Irf6* KO organoid line, with the majority of DEGs being downregulated relative to non-targeting control (189 for KO1, 187 for KO2) (**Fig. 6C-D, Table S5**). There was substantial congruence in DEGs between KO lines, with 111 shared down-regulated DEGs and 26 shared up-regulated DEGs (**Fig. 6D**). Likewise, GO pathways associated with *Irf6* KO DEGs were shared between the two KO organoid lines, including “epithelium development,” “cell death,” “cell adhesion,” and “regulation of immune system process” (**Fig. 6E**, **Table S5**). These pathways in *Irf6* KO organoids included substantial overlap with pathways altered in *Irf6* KO in M2C cell lines (**Fig. S1D**), increasing confidence in the association of Irf6 with epithelial homeostasis and immunity at baseline.

To determine which genes may be direct targets of Irf6, we compared our DEGs with IRF6 binding sites in ChIP-seq data from human keratinocytes (Botti et al., 2011). We saw 38 *Irf6*-associated DEGs were orthologs of genes from IRF6 ChIP-seq (**Table S6**), including three of the most highly down-regulated genes in both *Irf6* KO organoid lines: insulin-like growth factor binding protein 7 (*Igfbp7*), ADP ribosylation factor-like GTPase 4C (*Arl4c*), and pleckstrin homology-like domain family B member 2 (*Phldb2*) (**Fig. 6F**). *Phldb2* is associated with growth of cancer cells (Luo et al., 2022), and *Arl4c* plays roles in epithelial morphogenesis (Matsumoto et al., 2014), which is consistent with the reduced proliferation and size of *Irf6* KO organoids (**Fig. 5G-H**).

To further identify highest-confidence Irf6-regulated genes, we compared organoid DEGs to M2C cell line DEGs (**Fig. S1B-C**). 83 DEGs were shared between organoid KO and M2C KO lines (**Table S6**), including downregulation of genes associated with Wnt signaling (*Wnt10a*), regenerative stem cells (*Clu*), and maintenance of genome stability (*Zfp365*, also shared with IRF6 ChIP-seq) (**Fig. 6F**). Relatively few DEGs in *Irf6* KO organoids were upregulated, but some upregulated DEGs were indicative of increased differentiation: epithelial cell adhesion molecule (*Epcam*), mucin 13 (*Muc13*), and differentiation-promoting transcription factor caudal type homeobox 1 (*Cdx1*) (**Fig. 6F**).

To determine whether separate clusters could be identified within each experimental group, we performed unsupervised clustering of integrated single-cell data. We identified 14 clusters of differential gene expression (**Fig. 6G**). The clusters distinguished *Irf6* KO organoids from non-targeting controls, and identified distinct IFN-stimulated subsets (**Fig. S2B-C, Table S5**). Clusters 0-6 were distinguished by cell line and IFN treatment, confirming significant roles for Irf6 and IFN response in shaping transcriptional profile (**Fig. S2D-E**). Further analysis of clusters revealed that cell cycle phase was a major defining feature, with G1 phase represented in clusters 7-10, and G2M/S phases represented in clusters 0-6 (**Fig. 6H**). There was also a small cluster within untreated groups (**Fig. 6G**, cluster12) that expressed markers of secretory progenitor IECs, including master transcription factor *Atoh1*, Paneth cell-associated *Lyz1*, goblet cell-associated *Muc2*, and immunoglobulin transport receptor *Pigr* (**Fig. S2F**). This secretory progenitor cluster was predominantly found in *Irf6* KO1 organoids (**Fig. S2D**), suggesting a role of Irf6 in blocking secretory progenitor differentiation.

As an orthogonal test of differentiation status, we performed CytoTRACE analysis (Gulati et al., 2020), which examines transcriptional diversity to infer developmental potential. CytoTRACE analysis indicated that *Irf6* KO organoids had less differentiation potential at baseline (No IFN groups) compared to non-targeting control organoids (**Fig. 6I-J, S1G-H**). IFN treatments groups had further decreases in developmental potential within each group (**Fig. 6I-J**). Together, these data reveal a significant role for Irf6 in regulating the homeostatic transcriptome and developmental gene expression of primary IEC organoids.

Analysis of MNV genomes within the infected pool of cells indicated that most cells had a few counts (0-10 genomes/cell), and a minority of cells had 100-10,000 genomes/cell (**Fig. S2I-J**). These data indicate that most cells were not robustly infected, explaining why MNV was not a major driver of differential gene expression in this experiment. IFN-treated groups of all organoid lines had significantly fewer robustly-infected cells, and IFN-λ-treated *Irf6* KO2 cells were the only group that prevented any robustly-infected cells (**Fig. S2I-J**). However, there were no significant differences in MNV infection between cell lines. So, *Irf6*-dependent differences in antiviral control may be relatively nuanced in IEC organoids compared to epithelial homeostasis pathways.

### *Irf6* regulates ISG expression and ISRE activity in IEC organoids

To identify all IFN-regulated genes within organoid RNAseq data, we compared IFN-treated groups for each organoid line (NT, KO1, KO2) to their respective untreated controls. IFN types were combined for this analysis because there were minimal differences in clustering between IFN-β and IFN-λ treatments (**Fig. 6A** inset, **Fig. S2E**). We identified 162, 204, and 178 IFN-regulated genes for NT control, *Irf6* KO1, and *Irf6* KO2 organoids, respectively (**Fig. 7A, Table S5**). Many antiviral ISGs were similarly upregulated across all IFN-treated groups (e.g. *Ifih1, Tlr3*), but 20 ISGs were significantly higher in *Irf6* KO organoids relative to non-targeting controls (**Fig. 7B**). Some of these *Irf6*-dependent ISGs were favored within distinct IFN-stimulated KO subsets (**Fig. 6G**): *Muc3* and *Ifit1bl1* were preferentially stimulated in clusters 7 and 10 (G1 phase); *Ifitm3* and *Psmb8* were preferentially stimulated in clusters 0, 2, and 3 (G2/S phases) (**Fig. 7C**). Additionally, cluster 11 was unique to IFN-stimulated *Irf6* KO organoid lines and was distinguished by increased markers of apoptotic stress response (e.g. *Atf3*, *Atf4, Chac1*, **Figs. 8C, Table S5**), suggesting increased IFN-stimulated stress and cytotoxicity in *Irf6* KO organoids.

**Figure 7.**
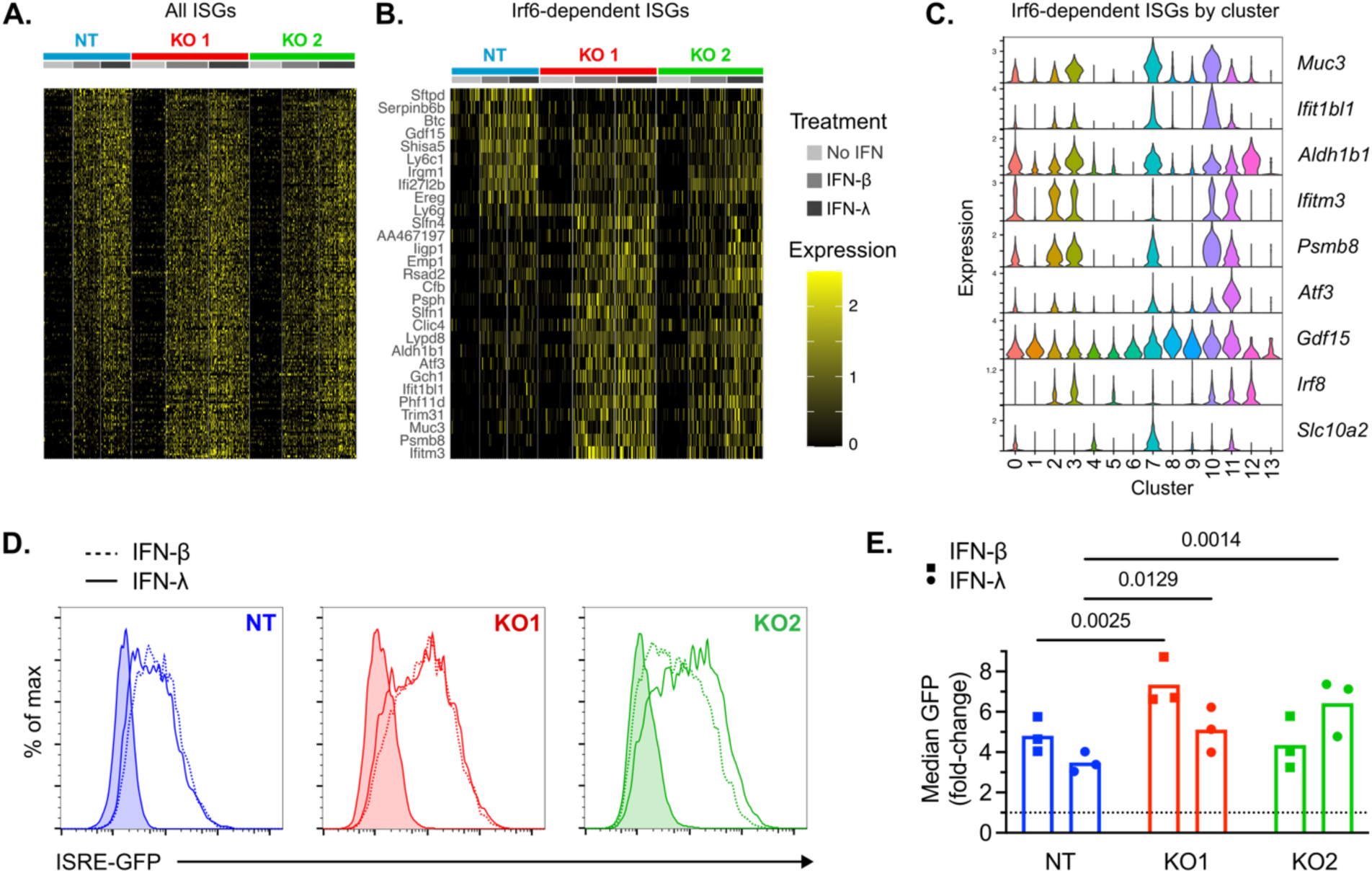
*Irf6* regulates the IFN response in primary IECs. **A-B.** Heatmaps of ISGs arranged by greater stimulation in non-targeting (top) to greater stimulation in *Irf6* KOs (bottom). **A.** all ISGs that are significantly increased by IFN treatment within at least one cell line **B.** ISGs that are significantly different between at least one KO line and non-targeting controls. **C.** Violin plots depicting expression of selected ISGs among clusters from figure 6G. **D-E.** Flow cytometry of Mx1-GFP expression 24hrs after treatment of indicated organoid lines with 10 ng/mL IFN-β (dashed lines) or 25 ng/mL IFN-λ (solid lines). **D.** Representative plots from three experimental replicates. **E.** Fold-change in median GFP expression of IFN-treated groups relative to their respective untreated controls. Data points represent replicates and significance was calculated using two-way ANOVA with Sidak multiple comparison correction.

**Figure 8.**
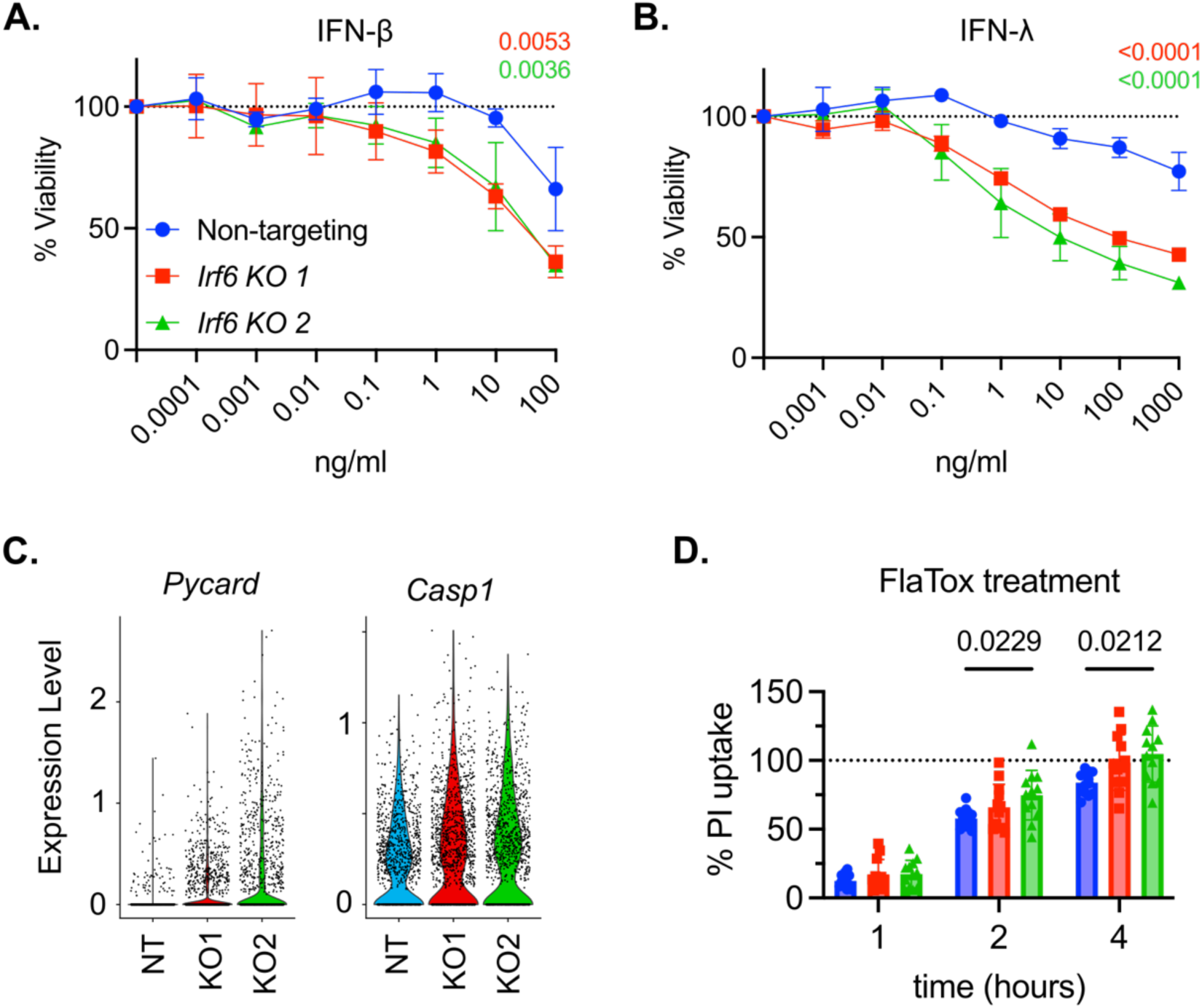
Increased innate immune cytotoxicity in Irf6-deficient IEC organoids. **A.** *Irf6* KO and non-targeting (NT) control organoids were treated with indicated concentrations of IFN-β (**A**) or IFN-λ (**B**) for 48hrs, and viable cells were quantified by ATPglo assay relative to no IFN treatment controls. Two independent replicates with statistical significance by two-way ANOVA. **C.** Gene expression for *Pycard* and *Casp1* from untreated cells in single-cell RNAseq data. **D.** Organoids were treated with propidium iodide (PI) viability stain in the presence or absence of FlaTox, and the percent of maximum PI fluorescence was measured relative to untreated control wells. Data points are combined from three independent experiments. Statistical significance by two-way ANOVA.

Further analysis of gene clusters identified some ISGs unique to subsets of each organoid line. For example, the bile acid cotransporter *Slc10a2* was preferentially stimulated in cluster 7 of KO2, but was more modestly stimulated in *Irf6* KO1 (**Fig. 7C**). Additionally, IFN-stimulated cluster 3 expressed secretory-lineage transcription factor *Atoh1* (**Fig. S2F**), suggesting ISGs in this cluster may be preferentially IFN-stimulated within secretory-lineage cells. These cluster 3 ISGs included aldehyde dehydrogenase (*Aldh1b1*) and *Irf8* (**Fig. 7C**). Furthermore, there was increased baseline expression of *Aldh1b1* and *Irf8* within untreated secretory progenitors (**Fig. 7C**, cluster 12), further linking these genes to IEC subsets. Together these data revealed clusters of IFN-stimulated response genes that correlated with *Irf6* expression, cell cycle phase, and secretory progenitor genes.

Differences in the ISG transcriptome of *Irf6* KO organoids may be related to differential IFN-stimulated activation of the ISRE promoter. The parental organoid line used to generate *Irf6* KOs was derived from the small intestine of an ISRE-GFP reporter mouse (Uccellini & García-Sastre, 2018).

Therefore, we used flow cytometry to quantify GFP reporter expression as an indicator of ISRE transactivation following 24 hours of treatment with either IFN-β or IFN-λ. All organoid lines had significantly higher GFP expression following IFN treatments, confirming the utility of this reporter gene (**Fig. 7D-E**). The median fold-increase in GFP fluorescence of *Irf6* KO1 organoids treated with either IFN type was significantly higher than non-targeting controls (**Fig. 7E**). *Irf6* KO2 organoids exhibited a preferential response to IFN-λ, with a significantly higher median fold-increase in GFP following treatment with IFN-λ, but not IFN-β (**Fig. 7E**). This IFN-λ phenotype of *Irf6* KO2 organoids was consistent with the result of preferential IFN-λ phenotype for *Irf6* in the viability CRISPR screen (**Fig. 2G-H**).

Together, these data support the conclusion that Irf6 dampens IFN responsiveness of IEC organoids.

### Increased innate immune cytotoxicity in Irf6-deficient IEC organoids

RNAseq data suggested that *Irf6* deficiency led to an increase in IFN-stimulated stress and cytotoxicity (cluster 11, **Fig. 6G, Table S5**). To quantify differences in IFN-stimulated cytotoxicity between IEC organoid lines, we treated cells with a titration of IFN-β or IFN-λ for 48 hours and quantified viability by ATPglo cell titer assay. Treatment with IFN-β concentrations below 1 ng/mL resulted in no appreciable change in viability, but 10 ng/mL IFN-β resulted in lower viability (63%) for each *Irf6* KO organoid line compared to non-targeting controls (95% viability) (**Fig. 8A**). Treatment with IFN-λ concentrations below 10 ng/mL resulted in no appreciable change in viability for non-targeting control cells, but concentrations as low as 0.1 ng/mL IFN-λ resulted in 85% viability for *Irf6* KO organoids (**Fig. 8B**). Viability decreased to 31-42% for *Irf6* KO organoids treated with 1000 ng/mL IFN-λ, whereas non-targeting controls were only reduced to 77% viability at this maximum concentration of IFN-λ (**Fig. 8B**). These data indicated that IFN treatment of IEC organoids results in a greater loss of viability in the absence of *Irf6*, particularly for IFN-λ treatment, which is usually not cytotoxic.

in addition to IFN-stimulated cytotoxicity, we hypothesized that inflammasomes may be more active in the absence of *Irf6*. We noted that the gene for inflammasome adaptor ASC (*Pycard*) was significantly upregulated in *Irf6* KO organoids at baseline (**Fig. 6F, 9C**). We also noted that the gene for inflammasome effector caspase 1 was significantly higher in KO organoids (**Fig. 8C**). Nlrc4 is an inflammasome protein expressed by mouse IECs that can be activated by NAIP proteins upon detection of cytosolic flagellin (Rauch et al., 2016, 2017). To test whether inflammasomes were differentially active in *Irf6*-deficient IECs, we quantified inflammasome-driven lysis of IEC organoid lines by stimulating the NAIP-NLRC4 inflammasome with agonist delivery to the cytosol (FlaTox) (von Moltke et al., 2012).

*Irf6* KO organoids exhibited significantly greater lysis following FlaTox addition compared to non-targeting control organoids (**Fig. 8D**). Taken together with IFN-stimulated cytotoxicity, these data indicate that IRF6-dependent gene expression programs can directly or indirectly moderate death of IECs following activation of innate immune pathways.

## Discussion

We set out to identify novel regulators of the IFN response in IECs through the use of complementary CRISPR screens, and discovered that targeting *Irf6* in M2C IEC cells (but not BV2 macrophages) led to increased IFN-stimulated protection against MNV infection (**Figs. 2-3**). We found that monoclonal isolates of *Irf6* KO M2C cells had a slower growth rate and decreased expression of epithelial development pathway genes (**Figs. 4-5**). Primary IECs express substantially more Irf6 than transformed M2C cells (**Fig. 5A**), and Irf6-deficient IEC organoids had a reproducible reduction in growth and differentiation genes as well as consistent alterations to ISG profile (**Figs. 6-7**). In particular, increased IFN-stimulated expression of stress genes (**Fig. 6**) was correlated with a greater cytotoxicity of IFN-treated *Irf6* KO IEC organoids (**Fig. 8A-B**). Thus, we have identified a novel role for IRF6 in shaping the biology of IECs at baseline, with attendant roles in regulating the response to IFN. This role extends to other immune pathways beyond IFN because we also found greater inflammasome-stimulated death in *Irf6*-deficient organoids (**Fig. 8C-D**).

IRF6 is known to be important for fidelity of orofacial development, and *Irf6* knockout mice are perinatal lethal with myriad developmental defects (Ingraham et al., 2006). IRF6 has been primarily studied as a lineage-defining transcription factor within the epidermis, and is known to promote expression of genes important for terminal differentiation of keratinocytes (Botti et al., 2011; Kousa et al., 2017; Oberbeck et al., 2019). Our findings suggest that IRF6 may play an analogous role in the development of IECs, with keratinocyte-specific transcriptional programs substituted for IEC-specific programs. Indeed, a recent study of human organoids identified *IRF6*-targeted cells to be significantly reduced in a pooled transcription factor screen (Lin et al., 2023), indicating an important role of IRF6 in human IECs. Additionally, a genome-wide association study of inflammatory bowel diseases identified a polymorphism within an *IRF6* intron that is associated with increased risk of disease (de Lange et al., 2017) and is associated with decreased expression of IRF6 transcripts. Thus, our observation of decreased developmental potential and increased cytotoxicity in *Irf6*-deficient IECs has potential implications for human disease. Future studies in IEC-specific conditional knockout mouse models will definitively test *Irf6* roles in development, immunity, and disease within intact tissues.

All IRF family transcription factors share a highly conserved DBD, and members of this transcription factor family with developmental roles could also participate in regulation of IFN-stimulated response genes. A dual role of IRF6 in development and immunity may be a beneficial strategy for shaping the immune response of epithelia to suit their physiological roles within tissues. Our data suggest that IRF6 restricts the IFN response of IECs, with increased stress and apoptosis pathway genes stimulated by IFN when IRF6 is absent (**Fig. 6**). This activity of IRF6 may be beneficial in reducing damage to epithelial cells during an active immune response in the intestine. Like the IFN response, inflammasome activation thresholds need to be properly balanced within IECs to balance capacity for pathogen clearance with cytotoxicity, and our data indicates a role for IRF6 in regulating this response threshold as well (**Fig. 8C-D**).

Increased expression of epithelial development genes such as *Muc2* in *Irf6* KO organoids suggests that secretory progenitor development may be limited by *Irf6* (**Fig. 5I**). Single-cell RNAseq data supports this possibility, with increased expression of secretory IEC transcription factor *Atoh1* and reduced expression of Notch ligand *Jag1* in *Irf6* KO organoids (**Fig. 6**). Organoid culture conditions used in this study maintain cells in high Wnt, which favors maintenance of stem cells. So, future studies testing organoid phenotypes under differentiation culture conditions that remove Wnt will be of interest.

The large, growth-arrested M2C cells observed within *Irf6* KO M2C cell isolation, and the significant increase in apoptosis pathway genes, suggests that these cells are experiencing greater genotoxic stress at baseline than non-targeting control cells. The selection pressure of genomic stress may have resulted in variable adaptations between KO lines. Alternatively, distinct phenotypes may result from the site targeted by each gRNA. *Irf6* gRNA 2 targets a sequence downstream of the DBD-encoding region, and it is possible that there is leaky expression of the resulting DBD-only truncated protein isoform. Such a DBD-only isoform would be predicted to act in a dominant-negative manner, with potential impacts extending to other IRF family members. This distinction between gRNA target sites may explain why we were unable to recover a homozygous knockout with *Irf6* gRNA 2 in IEC organoids as well as the substantially increased number of DEGs in the M2C cell line targeted with this gRNA.

We selected *Irf6* for further study from our screen, but *Irf2* was also found to play a substantial role in regulating the IFN-stimulated antiviral response in IEC cell lines (**Figs. 2-3**). IRF2 has been shown to bind ISRE elements and block IFN responses (Taki, 2002). Additional recent studies have implicated *Irf2* in IEC development, suggesting that it blocks IFN cytotoxicity of colonic stem cells (Minamide et al., 2020) or restricts differentiation into secretory lineages (Sato et al., 2020). It is intriguing to speculate that IRF6 and IRF2 may participate cooperatively or antagonistically in regulatory circuits related to IEC development and immunity. It will be interesting to define interaction between IRFs and other post-translational regulatory mechanisms for IRF6 in IECs. Regulation of IRF6 dimerization and nuclear translocation have been studied in keratinocytes, but it remains to be determined whether distinct mechanisms are at play in IECs. Further definition of these and other aspects of IRF6 regulation may have wide-ranging implications for intestinal homeostasis, immunity and disease.

## Methods

### Cell Culture

BV2 (macrophage) cells and HEK 293T cells (ATCC #CRL-3216) were maintained in DMEM (Gibco #11995065) with 5% fetal bovine serum (FBS), 1x penicillin/streptomycin/glutamine solution (Gibco #10378016), and 10 mM HEPES (HyClone #SH30237). M2C transformed colon epithelial cells (Padilla-Nash et al., 2012) were maintained with Advanced DMEM/F12 blend (Gibco #12634010) supplemented with 10% FBS, 1x penicillin/streptomycin/glutamine solution, and 10 mM HEPES. All cells and organoids were lifted and disrupted using trypsin/EDTA (Gibco #2500).

Organoids were generated, as previously described (Miyoshi & Stappenbeck, 2013), from the small intestine of a female MX1-GFP mouse (B6.Cg-*Mx1^tm1.1Agsa^*/J, Jackson Laboratory strain #033219). L-WRN cells (ATCC #CRL3276) were cultured for collection of conditioned supernatants containing Wnt3a, R spondin 3, and Noggin as previously described (Miyoshi & Stappenbeck, 2013). Organoid cultures were grown in Matrigel (Corning #354234) with 50% L-WRN conditioned media (CM) supplemented with 10 µm Y-27632 (MedChemExpress #HY10583) and 10 µm SB-431542 (MedChemExpress #HY10431).

### Mice

MX1-GFP mice were maintained in specific pathogen-free facilities at Oregon Health & Science University (OHSU). Animal protocols were approved by the Institutional Animal Care and Use Committee at OHSU (protocol #IP00000228) in accordance with standards provided in the *Animal Welfare Act*.

### Lentiviral production and cell transduction

Lentiviruses were produced from the following vectors: lentiCRISPRv2 hygro (Addgene #98291), pLenti CMV Blast empty (w263-1) (addgene #17486), and pCDH-MSCV CD300lf-T2A-GFP (gift from Dr. Craig Wilen). Insertion of gRNAs (**Table S1**) into the lentiCRISPRv2 hygro backbone was done as previously described (Shalem et al., 2014). CD300lf was cloned from a gene block (IDT) by amplifying with primers that included restriction site for XbaI and XhoI (**Table S1**). Vector backbone and CD300lf amplicon were restriction digested following manufacturer’s protocol. Fragments were gel purified and cloned using T4 DNA ligase. Chemically competent STBL3 *E. coli* were heat shock transformed with the ligated constructs and plated on ampicillin plates for selection. Resulting plasmid sequence was confirmed by sanger sequencing.

To produce the lentiviral particles, 293T cells were plated at 500,000 cells per well in a 6-well plate with 600 ng psPAX2, 300 ng pVSVg, 1000 ng lentiviral vector, 100 µl of Optimem (Gibco #31985-062) and 6 µl of Transit-LT1 (Mirus #MIR2300). Two days after transfection lentivirus was harvested and mixed with equal parts fresh media before overlaying on top of target cell lines. For transduction, BV2 cells were seeded at 20,000 cells per well and M2C cells were seeded at 1e4 cells per well in a 6-well plate. Two days after transduction lentivirus was removed and antibiotic selection media was added. After confirming death of untransduced control cells, transduced cell lines were cryogenically frozen in nine parts FBS one part DMSO.

Monoclonal cell lines were isolated by diluting polyclonal populations to 0.5 cells per 100 µl of media and 100 µl was plated in a 96-well plate. Wells were monitored for single cell colonies and CRISPR mutations were confirmed using NGS amplicon sequencing (Genewiz). Amplicons were PCR amplified for sequencing using Q5 polymerase with the corresponding primers (**Table S1**). Analysis of NGS sequencing data was done using CRISPResso2 (Clement et al., 2019).

Lentiviruses for the pooled CRISPR screen were produced as described above with equal proportions of all CRISPR/gRNA plasmids added to the transfection mix and the twelve wells of transfected 293T cells. The pooled lentiviral prep was used to infect 1000 cells per gRNA, at an MOI=0.5, as empirically determined for M2C and BV2 cells.

Lentiviral transduction of organoids was done after trypsinization to liberate from Matrigel and separate into single cells. The single cells were resuspended in a 1:1 mixture of 50% CM and lentiviral supernatant. The bottom of a 24-well plate was coated with 80 µl of Matrigel and solidified. The Cell/lentiviral mixture was then overlayed on top of the layer of Matrigel. Lentivirus was removed after 24hrs and replaced with 50% CM. Organoids were cultured and expanded for one week after transduction to allow for accumulation of the resistance gene within clonal organoids. After one week of culture, antibiotics were added to select for the transduced organoids. During selection, the surviving organoids were expanded. Selection antibiotics were removed for two days after each expansion to favor recovery after disruption of organoids and plating. Monoclonal organoids were generated by pipetting a single organoid into a new well and expanding. Mutations to the gRNA target site were determined as for cell lines above.

### Murine norovirus production, infection, and viability CRISPR screen

MNV was produced from molecular clones as previously described (Robinson et al., 2019). A chimeric strain CR6-VP1^CW3^ was used because it was shown to have the greatest lytic potential (Van Winkle et al., 2018), increasing the dynamic range of the survival screen. M2C and BV2 cell lines were seeded at 10,000 or 5,000 cells per well, respectively, in 96-well flat bottom black plates. At the time of plating, cells were treated with the indicated dosage of IFN-β (PBL #12405-1) or IFN-λ3 (PBL #12820-1). 24hrs after plating, cells were challenged with murine norovirus strain CR6-VP1^CW3^ at a MOI of either 50 for M2C cells or 10 for BV2 cells. 24 hours after infection, cell viability was quantified using the ATP-Glo™ Bioluminometric Cell Viability Assay (Biotium #30020-2) on a CLARIOstar plate reader.

For each CRISPR ko cell group, we calculated % viability compared to untreated controls, and calculated “% protection” attributed to IFN pretreatments by subtracting the viability of untreated conditions from viability of paired IFN-treatment conditions. We initially observed significant variance in % viability between CRISPR-transduced M2C-CD300lf cell lines after MNV infection (no IFN) that was independent of the specific gene targeted. To limit potential confounding effects of baseline variance in MNV susceptibility, and maximize the effect-size of IFN-treatment, we excluded poorly infected cells in which % viability following MNV infection was >50% in the absence of IFN pretreatment (**Table S2)**.

### Pooled CRISPR screen and FACS

Pooled populations of CRISPR cell lines were plated at 500,000 cells per 10 cm dish. After plating, BV2 cells were treated with 1 ng/ml IFN-β and M2C cells were treated with either 1 ng/ml IFN-β or 100 ng/ml IFN-λ. After 24hrs of IFN treatment, the cells were inoculated with MNV CR6-VP1^CW3^. BV2 cells were challenged with an MOI=10 and M2C cells were challenged with an MOI=100. After 8hrs the cells were lifted using trypsin. All the media and PBS used to wash the cells were collected and combined with the lifted cells to ensure any cells that died during infection were included in the sorting. Cells were stained with Zombie Aqua™ Fixable Viability Kit (Biolegend # 423102) and Fc receptors were blocked using the CD16/32 antibody (Biolegend #101302) for 20min on ice. Cells were washed with PBS and fixed in 2% paraformaldehyde for 20min at room temperature (RT). Cells were washed with PBS and permeabilized in PBS with 0.2% Triton X-100, 3%FBS, 1% normal goat serum (perm/block) for 30min RT. Cells were stored in perm/block at 4 degrees until both replicates had been collected. Immediately before sorting, cells were stained with a MNV NS1/2 polyclonal rabbit antibody (generous gift of Dr. Vernon Ward) for 30 minutes at room temperature. After washing two times, cells were stained with a goat anti-rabbit IgG antibody conjugated to Alexa 647 (ThermoFisher #A21244) in perm/block for 30 minutes at room temperature. Cells were washed twice with PBS 0.2% Triton X-100 and resuspended in FACS buffer for sorting.

The top 10% of cells stained with NS1/2 for each sample were sorted on the BD InFlux cell sorter for sequencing. DNA extraction was done using the Quick-DNA FFPE Miniprep (Zymo #D3067). Genome counts were determined through qPCR of the CRISPR insert (**Table S1**) and PCR amplification of the gRNA insert was done on 2000 genomes per sample using Q5® Hot Start High-Fidelity DNA Polymerase (NEB # M0493) with P5 and P7 primers that included the Genewiz partial adapter sequence (**Table S1**). Amplicons were purified with AMPure XP beads (Beckman Coulter # A63880) and submitted for amplicon sequencing (Genewiz). Analysis of gRNA sequences was done using MAGeCK (Li et al., 2014; B. Wang et al., 2019).

### Bulk RNA sequencing and analysis

RNA was extracted using the Zymo Quick-DNA/RNA Viral 96 Kit (ZymoResearch #D7023) from three M2C cell lines and three treatment groups each in triplicate experimental replicates (27 total samples). Quality of RNA samples were assessed using a TapeStation (Agilent) and mRNA sequencing libraries were prepared by the OHSU Massively Parallel Sequencing Shared Resource (MPSSR) using the TruSeq Stranded Poly(A)+ Library Prep Kit (Illumina). Barcoded libraries were pooled, paired-end sequencing was performed using the Illumina NovaSeq platform, reads were trimmed of adaptors, and reads were demultiplexed. Adaptor-trimmed and demultiplexed reads were mapped to the mouse genome (GRCm39) using the STAR aligner (Dobin et al., 2013), and mapping quality was evaluated using RSeQC (L. Wang et al., 2012) and MultiQC (Ewels et al., 2016). All samples had between 16 million and 27 million uniquely mapped reads with similar distributions across genomic features and uniform gene body coverage. Read counts per gene were determined using the featureCounts program (Liao et al., 2014), and differential expression analysis was performed using DEseq2 (Love et al., 2014), with each cell/treatment combination representing a different group in the study design (9 total comparison groups). PCA was performed on DEseq2 regularized logarithm (rlog)-transformed data. Heat maps were generated using either rlog-transformed raw counts or counts normalized to control samples (“Non-targeting” cells or “No IFN” treatment group), as indicated in figure legends. Heatmap clustering is based on Euclidean distance. Volcano plots were generated using the EnhancedVolcano program (https://github.com/kevinblighe/EnhancedVolcano).

### Single-cell RNA sequencing

For some experimental groups, clonal lentiCRISPR-transduced organoid lines (non-targeting, *Irf6* KO1, *Irf6* KO2) were further transduced with CD300lf using pCDH-MSCV CD300lf-T2A-GFP and transduction methods described above.

Each IEC organoid line was treated with 10 ng/mL IFN-β, 25 ng/mL IFN-λ, or media only. One group of replicate CD300lf-transduced organoids additionally received 9e5 PFU of MNV strain CR6-VP1^CW3^ at the same time as IFN treatments. 24 hours after treatments, single cells were prepared by incubation in trypsin/EDTA for 20 minutes, with pipetting every 5 minutes to disrupt organoids. The nine groups of cell lines (NT, KO1, KO2) and treatment conditions (no IFN, IFN-β, IFN-λ) were incubated with separate oligonucleotide-tagged antibodies (HTO) for multiplexing (Biolegend TotalSeq, A0301 - A0309). Groups were counted and pooled in equal abundance, with four separate pools of cells: two groups without CD300lf transduction or MNV infection, a CD300lf-transduced group without MNV infection, and a CD300lf-transduced group with MNV infection (**Fig. 6A**). Pools were submitted to the OHSU Gene Profiling Shared Resource for preparation of 10x chromium next GEM 3’ single cell gene expression v3 libraries and HTO libraries. Libraries from the four pools were prepared separately and sequenced on an Illumina NovaSeq by the OHSU MPSSR. Adaptor-trimmed and demultiplexed reads from the libraries of each pool were mapped with Cell Ranger Count v7.1.0 to the mouse genome (mm10-2020-A), with addition of MNV genome as a custom gene definition.

Gene counts from Cell Ranger were read into Seurat version 4.1.3 (Hao et al., 2021). Each pool was filtered for cells with less-than 10% mitochondrial reads, greater than 1000 genes, and greater than 5000 counts. Gene counts were normalized and variable features identified within each pool using the default parameters. Pools were integrated using FindIntegrationAnchors and IntegrateData functions (50 dimensions). HTO data was normalized using centered log-ratio (CLR) transformation, and groups were de-multiplexed using the HTODemux function (positive.quantile = 0.999). 12,151 demultiplexed singlets were clustered by gene expression using the following functions: ScaleData, RunPCA, FindNeighbors (dims = 1:15), FindClusters (resolution = 1), and RunUMAP (dims = 1:15). Experimental groups were identified by HTO, and differentially expressed genes between groups were identified using DESeq2 (Love et al., 2014) via the FindMarkers function. DEGs were defined as having a greater than 1.5 fold-change and adjusted p-value < 0.05. Marker genes for clusters were identified by Wilcoxon Rank Sum test using the FindAllMarkers function (min.pct = 0.25). Cell cycle phase was determined using the CellCycleScoring() function.

### Quantitative PCR

RNA was extracted using the Zymo Quick-RNA Viral Kit (ZymoResearch #R1035). DNA contamination was removed using the Turbo DNAfree kit (ThermoFisher #AM1907). cDNA was generated with the ImPromII reverse transcription system (Promega #A3800). Quantitative PCR was performed using PerfeCTa qPCR FasMix II (QuantaBio #95119) and the pre-designed primer and probe assays from Integrated DNA Technologies (IDT) (**Table S1**). Absolute copy number was determined by comparing Ct values to a standard curve generated using DNA of known copy number encoding the target sequences. Samples are graphed as absolute copy number of the indicated target divided by the absolute copy number of the housekeeping gene, *Rps29*, with log-transformation and normalization as indicated in figure legends.

### Western blot

Two days after plating, organoids were dissociated from Matrigel using trypsin/EDTA (Gibco #2500), washed with PBS, and lysed in RIPA buffer (NaCl 150mM, Tris-HCl 50mM [pH 8.0], sodium deoxycholate 0.5%, and SDS 0.1%) supplemented with cOmplete mini, EDTA-free protease inhibitor cocktail (Sigma #4693159001). Each sample was mixed with Bolt LDX buffer (ThermoFisher), Bolt reducing agent (ThermoFisher), and incubated at 70C for 10 min. Samples were run on a 12% Bis-Tris Bolt Mini protein gel (ThermoFisher) and transferred to a PVDF membrane using Bolt transfer buffer (ThermoFisher). IRF6 antibody (BioLegend #674502) was diluted 1:500 and the secondary antibody goat anti-mouse conjugated to horseradish peroxidase (ThermoFisher #62-6720) was diluted 1:5000.

### Fluorescence *in situ* hybridization

FISH was performed using the Advanced Cell Diagnostics (Newark, CA) manual RNAscope assay following manufacturer protocol from FFPE tissue sections. Probes specific for *Mus musculus* genes *Irf6* (ACD, #462931) and *Ifit1* (ACD, #500071-C2) were purchased from Advanced Cell Diagnostics. Slides were counter-stained with DAPI and mounted with ProLong Gold antifade reagent (ThermoFisher). Fluorescent micrographs were captured using a Zeiss ApoTome2 on an Axio Imager, with a Zeiss AxioCam 506 (Zeiss) detector.

### FlaTox inflammasome assay

Organoids were seeded into 5μL Matrigel domes in 96-well plates, at least 3 wells per treatment. After 2-3 days, organoids were treated with FlaTox (comprised of flagellin from *V. parahemolyticus* (16μg/mL) and protective antigen (1μg/mL)) and propidium iodide (1:100 dilution) in complete media. Absorbance was measured using on a CLARIOstar plate reader each hour following treatment. Absorbance readings were first normalized to untreated controls (0%) and then normalized to maximum PI uptake in replicate wells for each organoid line treated with 1% Triton-X (100%).

### Statistical Analyses

Sample size estimation was performed based on historical data. Data were analyzed with Prism software (GraphPad Prism Software), with specified tests as noted in the figure legends.

### Data availability

RNA sequencing data obtained in this study have been deposited in the NCBI gene expression omnibus (GEO) under series number GSE245972.

## Acknowledgments

The authors would like to thank the following OHSU core facilities: Integrated Genomics Laboratory, Massively Parallel Sequencing Shared Resource, Advanced Light Microscopy Core, and Flow Cytometry Core. T.J.N. and A.P.W. were supported NIH grant R01-AI130055 and the Medical Research Foundation of Oregon. T.J.N. was additionally supported the Kenneth Rainin Foundation (Grant #20220034), and OHSU school of medicine. The funders had no role in study design, data collection and interpretation, or the decision to submit the work for publication.

---

**Figure S1.**
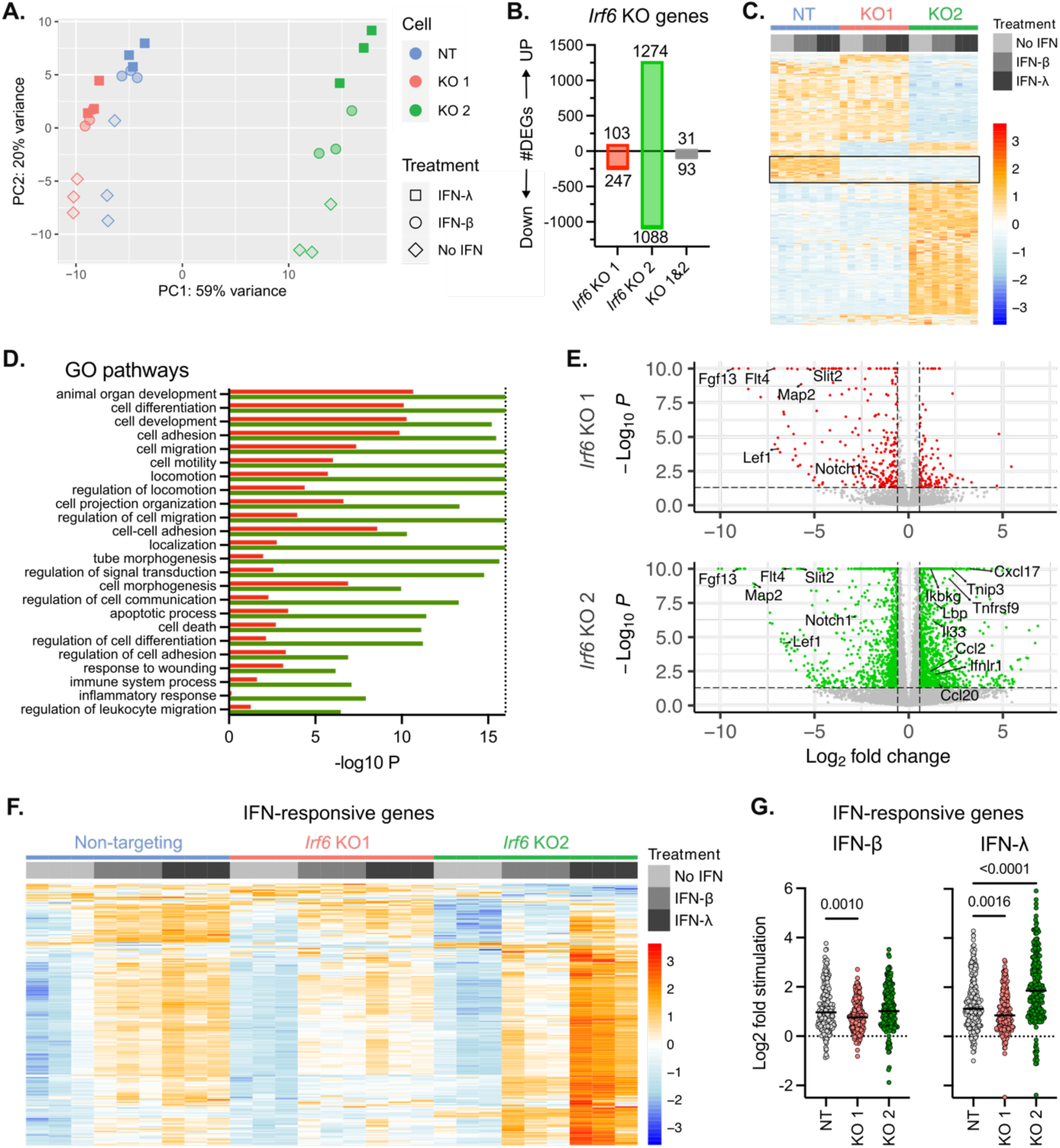
*Irf6* KO alters baseline and IFN stimulated gene expression. Related to figure 4. RNA sequencing of *Irf6* KO M2C cells and non-targeting controls with or without 24hrs of IFN treatment. All data is from three experimental replicates. **A.** Principal component plot. **B-E.** Differential gene expression of *Irf6* KO cells relative to non-targeting controls within the untreated (‘No IFN’) groups **B.** Differentially expressed genes (adjusted p-value < 0.05 and fold-change > 1.5). **C.** Heatmap of DEGs from **B**, with data scaled and centered by row. Box highlights DEGs down-regulated in both KO cell lines. **D.** Selected GO pathways significantly associated with DEGs from *Irf6* KO1 (red bars) and *Irf6* KO2 (green bars) cell lines. **E.** Volcano plot of all genes; dotted line represents cut-off for DEGs. **F.** Heatmap of IFN-responsive genes, including genes differentially expressed in at least one IFN-treated group compared to respective no IFN control. Data scaled and centered by row. **G.** Log2 fold-change of genes from **F**. P-values calculated by Kruskal-Wallis test.

**Figure S2.**
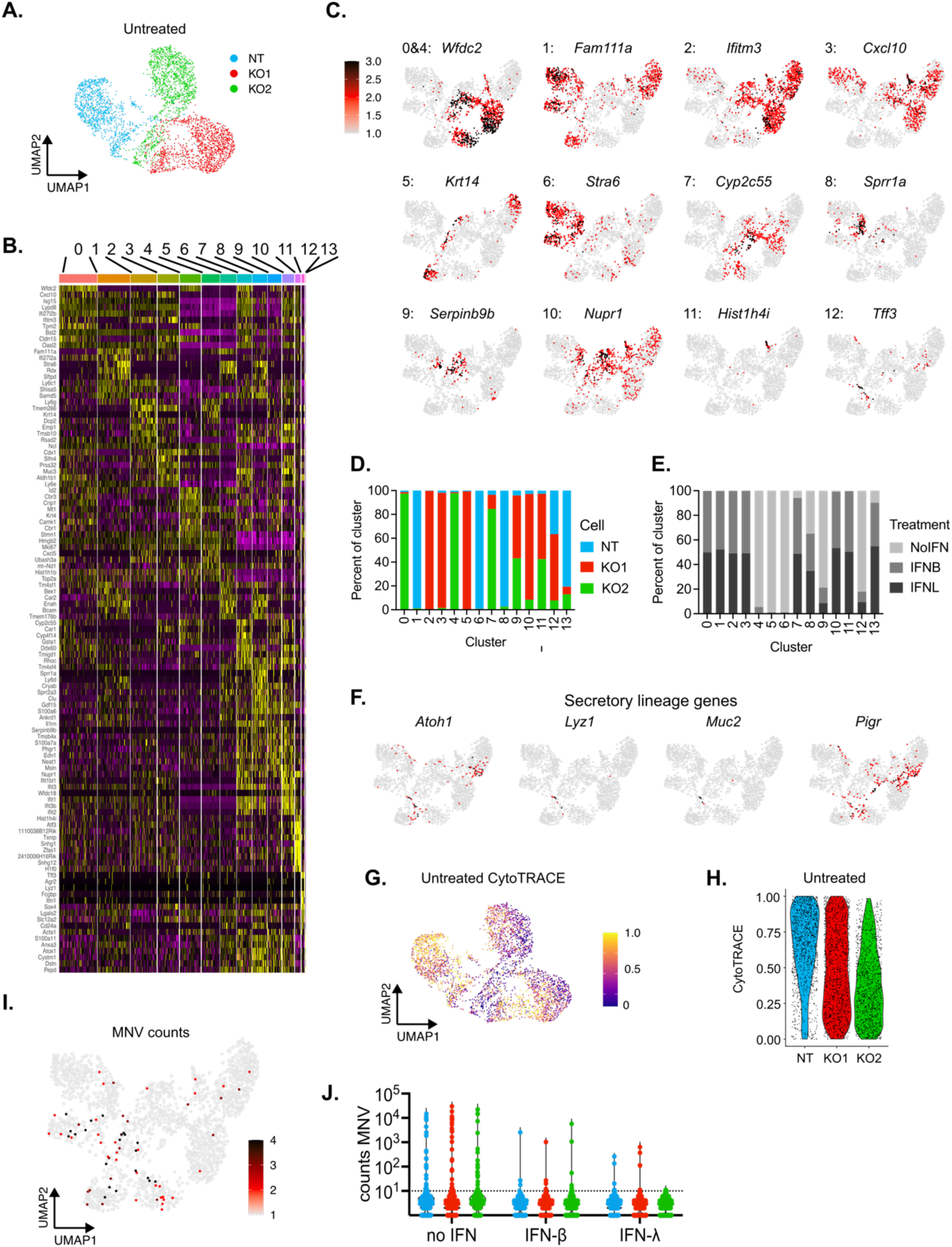
Related to figure 6. **A.** UMAP clustering of untreated cells only. **B.** Heatmap of 10 genes for each cluster (Fig. 6G) with the greatest fold-change relative to all other clusters. **C.** Feature plots depicting the distribution of expression for the top gene for each cluster. **D-E.** Distribution within each cluster of cell (**D**) or treatment (**E**) identities. **F.** Feature plots depicting selected genes enriched in cluster 12. **G-H.** CytoTRACE analysis of differentiation on untreated cells only. Higher CytoTRACE score indicates more stem-like cells. **I**. Feature plot of MNV genome count distribution among all groups. **J.** Violin plot of MNV counts within the MNV-infected pool only.

